# Relationship between Parathyroid Hormone Levels and Hazards of Fracture, Vascular Events and Death in Stage 3 and 4 Chronic Kidney Disease

**DOI:** 10.1101/480848

**Authors:** Sinong Geng, Zhaobin Kuang, Peggy L. Peissig, David Page, Laura Maursetter, Karen E. Hansen

## Abstract

**Background and Objectives:** Chronic kidney disease (CKD) affects ∼20% of older adults and secondary hyperparathyroidism (HPT) is a common condition in these patients. Studies have linked HPT to a greater risk of fractures, vascular events and mortality. However, the optimal parathyroid hormone (PTH) level needed to minimize these events remains uncertain.

**Design, setting, participants and measurements:** We assessed relationships between baseline serum PTH levels and the subsequent 10-year probability of clinical fractures, vascular events and death in stage 3 and 4 CKD patients. We used Marshfield Clinic Health System electronic health records to analyze data from adult CKD patients spanning from 1985 to 2013. We required ≥2 PTH measurements at baseline and used ICD-9 codes to identify medical conditions, fractures, vascular events and death. In multivariate models, we assessed relationships between serum PTH and the three clinical outcomes, controlling for age, gender, co-morbidities and osteoporosis medication.

**Results:** 7594 subjects had a mean age of 68±13 years and 55% were women. Fractures, vascular events and death occurred in 19%, 60% and 29% of the cohort, respectively. In multivariate models including the whole cohort regardless of PTH assay, the probability of fracture, vascular events and death were minimized at a PTH of 23, 50 and 50 pg/mL. Below these cutpoints, the probability of fractures and death dramatically increased. When confining the analysis to patients measured using a 2^nd^ generation PTH assay (n=5108), the hazards of fracture, vascular events and death were minimized at a PTH of zero, 60 and 58 pg/mL. Any of these clinical outcomes was minimized at a baseline PTH of 58 pg/mL.

**Conclusions:** Our study suggests that parathyroid hormone levels around 60 pg/mL might reduce the risk of fractures, vascular events and death in CKD patients. Additional epidemiologic studies and randomized clinical trials are needed to confirm these findings.

## Introduction

CKD affects one in five older adults (1) and 14% of the general population (2). CKD causes more deaths per year than breast or prostate cancer (2). Patients >65 years old with CKD have a higher prevalence of cardiovascular disease (70% vs. 35%) compared to individuals of the same age without CKD (2). Medicare spends almost one-fifth of its budget on CKD patient care, due to excess vascular events, fractures, infections and subsequent premature death (3) compared to patients without CKD. Secondary hyperparathyroidism (HPT) affects nearly all stage 5 (dialysis) CKD patients and over half of stage 3-4 CKD patients (4-6). The pathogenesis of hyperparathyroidism is well established (7). Declining renal 1,25(OH)_2_D synthesis leads to reduced intestinal calcium absorption and lower ionized serum calcium levels. To maintain normocalcemia, the parathyroid glands released parathyroid hormone (PTH), which upregulates renal 1,25(OH)_2_D synthesis to increase calcium absorption, and stimulates osteoclastic bone resorption to liberate skeletal calcium into the bloodstream.

Most cross-sectional studies in CKD patients have found an inverse relationship between PTH and bone mineral density (BMD) (5, 8-11), suggesting that HPT contributes to fractures. Researchers have also found that high PTH was associated with vascular calcification (12) and vascular events (13) in CKD patients. From these studies, we hypothesized that treating HPT in stage 3-4 CKD patients might increase BMD and reduce the risk of fractures, vascular events and death. We found weak evidence suggested that treating HPT improved bone mineral density (2 trials, 62 subjects) (14, 15) and no clinical trials demonstrating that treatment of HTP reduced vascular events or death in stage 3-4 CKD patients.

Based on expert opinion and observational studies, the National Kidney Foundation published guidelines on how to manage HPT. The 2003 guidelines (16) suggested specific PTH targets to achieve, depending on the stage of kidney disease (**Appendix Table 1**). However, the 2009 National Kidney Foundation guidelines (7) acknowledged that target PTH levels were based on expert opinion, rather than randomized clinical trials. Because vitamin D analogs can raise serum calcium and phosphorus and potentially promote vascular calcification, the 2017 National Kidney Foundation guidelines (17) reserved treatment for patients with severe and progressive hyperparathyroidism; specific PTH targets were not recommended.

**Table 1:** Subject Characteristics

| Table 1: Subject Characteristics |  |  |
| --- | --- | --- |
|  |  | All Subjects<br>n=7594 |
| Clinical | Age, years | 68 ± 13 |
|  | Female Gender | 4176 (55%) |
|  | Tobacco use | 4129 (54%) |
|  | Duration of care at Marshfield Clinic | 24 ± 15 years |
| Prevalence of Comorbidities | Coronary artery or peripheral vascular disease | 4981 (66%) |
|  | Diabetes | 3875 (51%) |
|  | Hypertension | 6793 (89%) |
|  | Hyperlipidemia | 3606 (47%) |
|  | Obesity | 3141 (41%) |
| Labs | GFR, mL/minute | 39 ± 16 |
|  | PTH, pg/mL | 98 ± 91 |
| Medication | Bisphosphonates | 100 (1.3%) |
|  | Denosumab or teriparatide | 3 (0.04%) |
|  | Estrogen, raloxifene, testosterone | 62 (0.8%) |
| Outcomes | Fractures | 1408 (19%) |
|  | Vascular Events | 4533 (60%) |
|  | Death | 2196 (29%) |

In summary, observational studies have linked HPT to lower BMD, excess fractures and excess vascular events that together would shorten lifespan. However, potential harm from use of vitamin D analogs, and lack of clinical trials proving the benefits of such therapy, have led to uncertainty about whether to treat HPT. We utilized the Marshfield Clinic Health System (MCHS) data repository to evaluate relationships between serum parathyroid hormone levels and clinical fractures, vascular events, and mortality in patients with stage 3 and 4 chronic kidney disease. Our objective was to identify the PTH level that minimized these three adverse clinical outcomes. We hypothesized that higher parathyroid hormone levels would be associated with greater probability of fractures, vascular events and death, compared to lower parathyroid hormone levels.

## Materials and Methods

We utilized electronic health records of patients within the MCHS data repository from a collection period spanning ∼30 years (1985 to 2013). The database includes patients’ demographic characteristics, medical conditions, laboratory results, fractures, imaging results and medication use. We included patients who were 1) ≥21 years old, 2) had a mean GFR <60 mL/minute during the baseline year based on the average of at least 2 measures, 3) had at least two parathyroid hormone levels measured during the baseline year during outpatient clinic visits, and 4) subsequently received medical care at Marshfield Clinic for at least 2 years, prior to November 2013. We excluded patients on dialysis and patients with primary hyperparathyroidism. We further excluded patients who had undergone parathyroidectomy, a procedure which is largely restricted to dialysis patients and is associated with increased mortality (18).

At least two ICD-9 coded clinic encounters within a 24-month interval were required to denote the presence of diabetes, hypertension or hyperlipidemia, as described elsewhere (19, 20). We used ICD-9 codes designating specific skeletal fracture sites known to reflect osteoporotic fracture including hip (ICD-9 820-822), wrist (ICD-9 813), humerus (ICD-9 812) and vertebral (ICD-9 805) fractures. We also used ICD-9 codes to denote vascular events. We required that at least two clinical encounters noted the ICD-9 code prior to counting the vascular event, since “ruling out” such an event was often used to support the need for imaging (e.g. head CT scan to rule out stroke).

## Statistical Analysis

We analyzed predictors of three clinical outcomes (fracture, vascular events, death) occurring after parathyroid hormone measurement, using the mean of two or more baseline PTH values. First, we used survival tree models (21) to identify the PTH level at which the probability of each clinical outcome was minimized. In the full survival tree model, we used all covariates while in the second or reduced model we employed the mean PTH level alone. After ten-fold cross validation, we identified the PTH level at which the probability of each outcome was minimized, then we divided patients into groups according to whether their PTH level was above or below that level. We used Cox proportional hazard ratios and Kaplan-Meier survival curves to evaluate the risks of each clinical outcome in patients with PTH below and above the “optimal” level identified by survival tree models. Our multivariate models controlled for age, gender, tobacco use, diabetes, hypertension, hyperlipidemia, vascular disease, obesity, GFR, bisphosphonate and other bone active medications (22-24).

As in other medical centers, the PTH assay used at the Marshfield Clinic Laboratory changed several times over the span of 30 years. First generation PTH assays measured PTH fragments that accumulated in renal failure but were devoid of biologic activity (25). Third generation PTH assays did not detect PTH7-84 fragments (25). Currently, the 2^nd^ generation assay is used in the vast majority of medical centers (26). To make our study results most relevant to current clinical practice, we performed a sensitivity analysis in which the three clinical outcomes were analyzed by excluding subjects whose PTH was measured using a 1st or 3^rd^ generation PTH assay. We performed a further sub-analysis to assess whether the optimal PTH level differed between patients with stage 3 and stage 4 CKD. We used version 3.2.3 of “R” (The R Project for Statistical Computing, http://www.r-project.org) for all statistical analyses.

## Results

We identified 7594 eligible subjects (55% female) with a mean ± standard deviation age of 68 ± 13 years (**Table 1**). Patients received care at MCHS for 24 ± 15 years. Among the cohort, 19% sustained clinical fractures, 60% experienced vascular events and 29% died. We did not include race, body mass index, denosumab or teriparatide as co-variates due to lack of records regarding these data (race, body mass index) or too few individuals with the characteristic (use of denosumab or teriparatide). The 2010 United States Census estimated that ∼19,000 people lived in Marshfield, Wisconsin and the vast majority (∼95%) were Caucasian, suggesting that our cohort was largely Caucasian (27). We had no missing data for age, gender, diabetes, hypertension, hyperlipidemia, coronary artery disease, peripheral vascular disease, bisphosphonate, estrogen or testosterone therapy.

### Analysis of all subjects regardless of PTH assay

Age, gender, vascular disease, obesity, GFR and PTH were significant co-variates in the multiple variable model predicting fracture (**Table 2**). The PTH value at which the probability of fracture was minimized equaled 23 pg/mL (**Figure 1**). Below this level, the 10-year probability of fracture dramatically increased. Surprisingly, at PTH values >23 pg/ml, the probability of fracture increased only modestly. In the multiple variable model predicting vascular events, age, gender, diabetes, hypertension, hyperlipidemia, obesity, GFR and PTH were significant co-variates (**Figure 1**, **Table 2**). The PTH value at which the probability of vascular events was minimized equaled 50 pg/mL (**Table 2, Figure 1**). Finally, age, gender, tobacco, vascular disease, diabetes, hypertension, hyperlipidemia, obesity, GFR and PTH were each significant co-variates in the multiple variable models predicting death (**Table 2).** The PTH level associated with highest probability of survival was 50 pg/mL. Just as with fractures, the probability of death increased sharply with lower PTH levels (**Figure 1**).

**Table 2:** Multiple Co-Variate Models Predicting Hazard Ratios for Fracture, Vascular Events and Death in 7594 Subjects, with PTH measured using any assay

|  |  |  |  |  |  |  |  |
| --- | --- | --- | --- | --- | --- | --- | --- |
|  |  | Fracture |  | Vascular Events |  | Death |  |
| Covariates |  | HR (95% CI) | p-value | HR (95% CI) | p-value | HR (95%CI) | p-value |
| Clinical | Age ≤69 years | 0.32 (0.27, 0.38) | <0.001 | 0.34 (0.31, 0.38) | <0.001 | 0.18 (0.16, 0.21) | <0.001 |
|  | Age 70-79 | 0.35 (0.30, 0.42) | <0.001 | 0.48 (0.43, 0.53) | <0.001 | 0.31 (0.27, 0.35) | <0.001 |
|  | Age 80-89 | 0.46 (0.40, 0.53) | <0.001 | 0.63 (0.58, 0.68) | <0.001 | 0.44 (0.40, 0.49) | <0.001 |
|  | Age ≥90 years | referent |  | referent |  | referent |  |
|  | Female Gender | 1.65 (1.47, 1.85) | <0.001 | 0.77 (0.72, 0.81) | <0.001 | 0.73 (0.67, 0.80) | <0.001 |
|  | Tobacco Use | 0.89 (0.81, 1.00) | 0.058 | 1.04 (0.98, 1.11) | 0.174 | 0.77 (0.71, 0.84) | <0.001 |
| Comorbidities | Vascular disease | 1.62 (1.42, 1.85) | <0.001 | - | - | 2.14 (1.88, 2.43) | <0.001 |
|  | Diabetes | 1.03 (0.92, 1.15) | 0.567 | 1.40 (1.32, 1.49) | <0.001 | 1.24 (1.14, 1.36) | <0.001 |
|  | Hypertension | 0.98 (0.79, 1.20) | 0.823 | 1.18 (1.04, 1.33) | 0.010 | 0.65 (0.56, 0.75) | <0.001 |
|  | Hyperlipidemia | 0.93 (0.83, 1.04) | 0.198 | 1.10 (1.04, 1.17) | 0.008 | 0.82 (0.75, 0.8964) | <0.001 |
|  | Obesity | 0.87 (0.77, 0.98) | 0.018 | 1.11 (1.04, 1.18) | 0.002 | 0.85 (0.77, 0.93) | <0.001 |
| Labs | GFR, mL/minute | 1.01 (1.01, 1.01) | <0.001 | 0.99 (0.99, 1.00) | <0.001 | 0.97 (0.96, 0.9706) | <0.001 |
|  | PTH > cutpoint<sup>a</sup> | 1.00 (1.00, 1.00) | 0.0376 | 1.00 (1.00, 1.00) | <0.001 | 1.00 (1.00, 1.00) | <0.001 |
|  | PTH < cutpoint<sup>a</sup> | 0.97 (0.96, 0.98) | <0.001 | 1.00 (0.99, 1.00) | 0.621 | 0.98 (0.98, 0.99) | <0.001 |
| Meds | Biophosphonates | 1.27 (0.86, 1.87) | 0.231 | 0.83 (0.63, 1.08) | 0.169 | 0.82 (0.55, 1.23) | 0.293 |
|  | Other<sup>b</sup> | 0.89 (0.49, 1.61) | 0.695 | 1.02 (0.74, 1.41) | 0.901 | 0.84 (0.50, 1.43) | 0.600 |

<sup>a</sup>PTH cutpoint = 23 pg/mL for fractures, 50 pg/mL for vascular events, and 50 pg/mL for death
<sup>b</sup>Other medications included estrogen, raloxifene and testosterone

**Figure 1:**
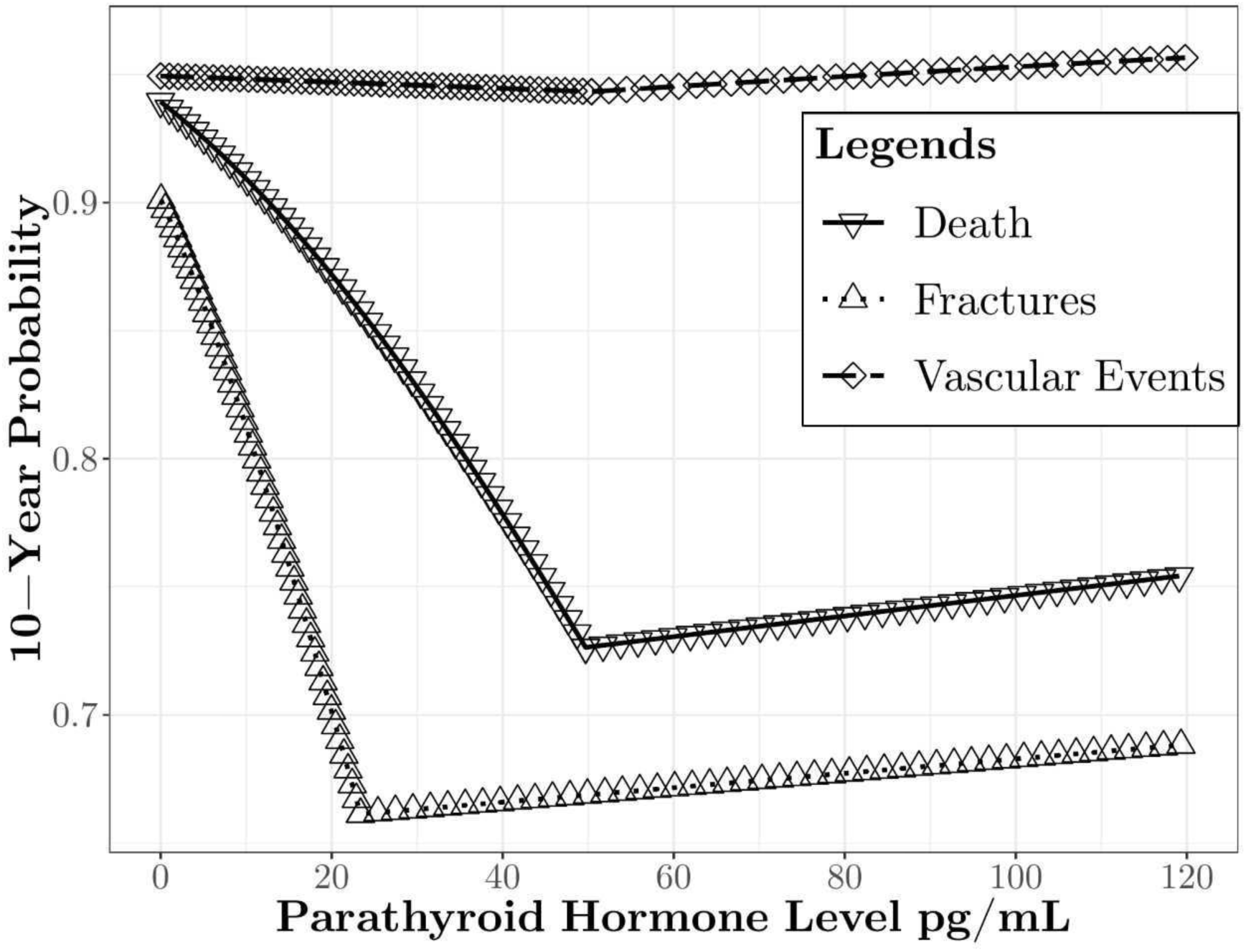
Ten-Year Probability of Fractures, Vascular Events and Death based on Baseline Parathyroid Hormone Levels.

In a 5-year survival analysis, we applied a logistic regression to the data that included patients who were alive or deceased 5 years after the diagnosis of stage 3-4 CKD. With a 10-fold cross validation, we drew the receiver operating characteristic (ROC) curve for prediction that had an area under the curve (AUC) value of 0.784. We used age, gender, tobacco use, vascular disease, diabetes, hypertension, hyperlipidemia, obesity, GFR, PTH, bisphosphonates, and other drugs (estrogen, raloxifene, testosterone) as covariates predicting death.

### Analysis restricted to subjects measured using second generation PTH assay

Among 5108 individuals who had baseline PTH levels measured using a second generation assay, age, gender, vascular disease, osteoporosis therapy and PTH were significant co-variates in the multiple variable model predicting fracture (**Table 3**). Subsequent models suggested that a PTH level of 101 pg/mL was associated with lowest subsequent risk of fracture. However, a comparison of patients with PTH values below and above this level indicated no significant difference in the risk of fracture (**Figure 2**). In fact, Figure 2 demonstrates that a PTH level of zero was associated with the lowest probability of subsequent fracture.

**Table 3:** Multiple Co-Variate Models Predicting Hazard Ratios for Fracture, Vascular Events and Death in 5108 Patients with 2^nd^ Generation PTH Values Above and Below Thresholds

|  |  |  |  |  |  |  |  |
| --- | --- | --- | --- | --- | --- | --- | --- |
|  |  | Fracture |  | Vascular Events |  | Death |  |
| Covariates |  | HR (95% CI) | p-value | HR (95% CI) | p-value | HR (95%CI) | p-value |
| Clinical | Age ≤59 years | 0.47 (0.36, 0.63) | <0.001 | 0.44 (0.39, 0.49) | <0.001 | 0.30 (0.26, 0.34) | <0.001 |
|  | Age 60-69 | 0.54 (0.42, 0.69) | <0.001 | 0.56 (0.50, 0.62) | <0.001 | 0.40 (0.36, 0.45) | <0.001 |
|  | Age 70-79 | 0.75 (0.60, 0.92) | 0.006 | 0.70 (0.63, 0.77) | <0.001 | 0.58 (0.52, 0.64) | <0.001 |
|  | Age ≥80 years | referent |  | referent |  | referent |  |
|  | Male Gender | 0.69 (0.58, 0.84) | <0.001 | 1.28 (1.18, 1.38) | <0.001 | 1.26 (1.16, 1.38) | <0.001 |
|  | Tobacco Use | 0.93 (0.79, 1.11) | 0.427 | 1.07 (0.99, 1.16) | 0.069 | 1.01 (0.93, 1.10) | 0.752 |
| Comorbidities | Vascular disease | 1.80 (1.44, 2.26) | <0.001 | - | - | 1.61 (1.44, 1.79) | <0.001 |
|  | Diabetes | 1.18 (1.00, 1.40) | 0.055 | 1.29 (1.19, 1.39) | <0.001 | 1.23 (1.13, 1.33) | <0.001 |
|  | Hypertension | 1.28 (0.84, 1.94) | 0.249 | 1.05 (0.89, 1.23) | 0.596 | 0.75 (0.64, 0.88) | <0.001 |
|  | Hyperlipidemia | 0.99 (0.83, 1.17) | 0.887 | 0.99 (0.93, 1.07) | 0.879 | 0.80 (0.74, 0.86) | <0.001 |
|  | Obesity | 0.89 (0.75, 1.07) | 0.212 | 1.12 (1.04, 1.21) | 0.003 | 0.91 (0.84, 0.99) | 0.029 |
| Labs | GFR, mL/minute | 1.00 (0.99, 1.00) | 0.268 | 0.99 (0.99, 0.99) | <0.001 | 0.98 (0.98, 0.98) | <0.001 |
|  | PTH < cutpoint<sup>a</sup> | 0.95 (0.78, 1.15)<sup>b</sup> | 0.605 | 1.11 (1.00, 1.23) | 0.058 | 1.05 (0.89, 1.25) | 0.546 |
|  | PTH > cutpoint<sup>a</sup> | 1.16 (0.93, 1.45)<sup>b</sup> | 0.195 | 1.18 (1.09, 1.27) | <0.001 | 1.19 (1.09, 1.30) | <0.001 |
| Meds | Biophosphonates | 1.65 (1.35, 2.01) | <0.001 | 1.00 (0.90, 1.11) | 0.988 | 1.09 (0.98, 1.22) | 0.126 |
|  | Other<sup>c</sup> | 1.50 (1.22, 1.85) | <0.001 | 0.93 (0.83, 1.03) | 0.176 | 0.89 (0.79, 1.01) | 0.068 |

<sup>a</sup>PTH cutpoint = 101 pg/mL for fractures, 69 pg/mL for vascular events and 58 pg/mL for death
<sup>b</sup>In the fracture model using PTH as a continuous variable, PTH remained a significant independent risk factor (p=0.042).
<sup>c</sup>Other medications included estrogen, raloxifene and testosterone

**Figure 2:**
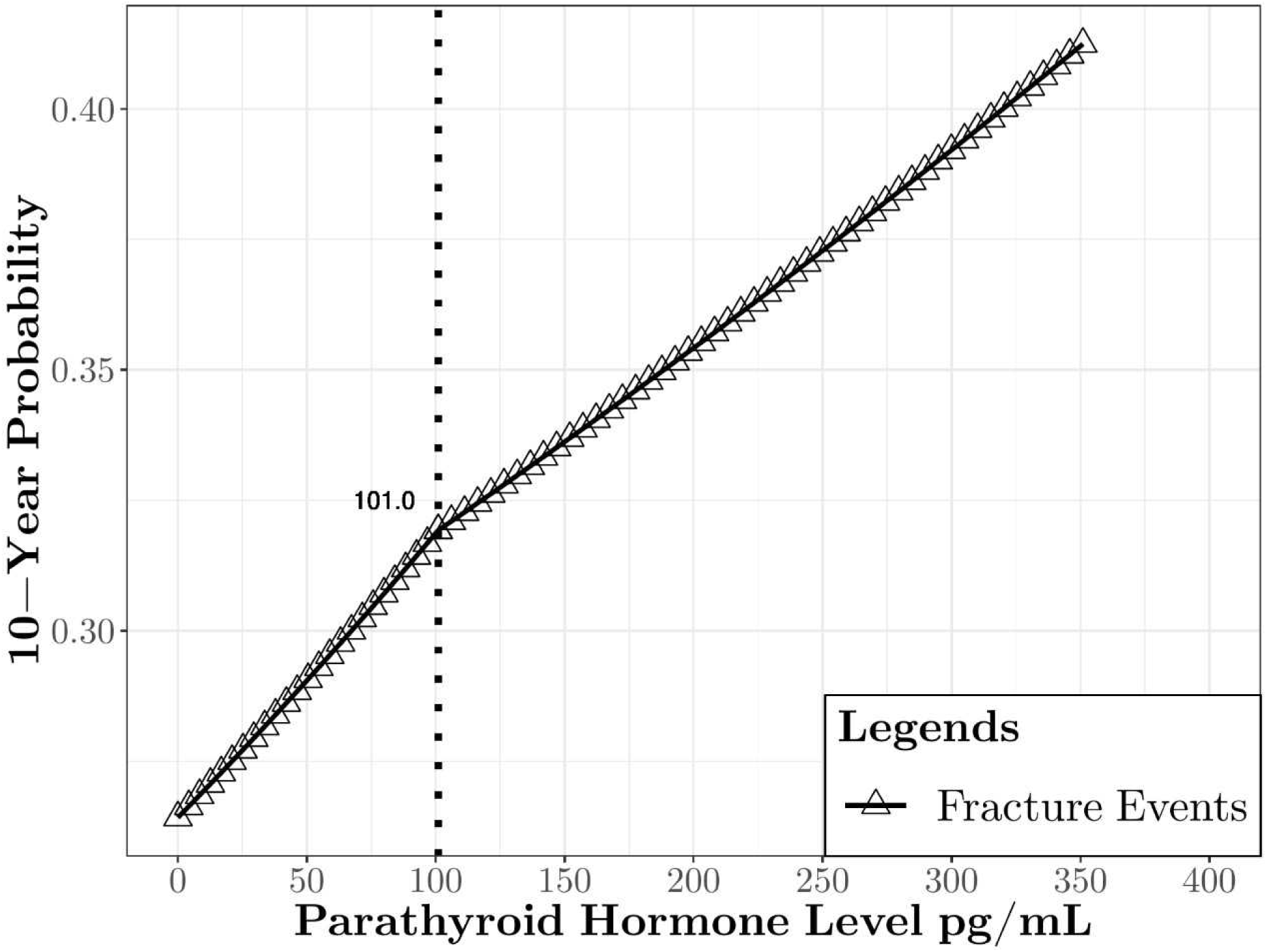
Ten-Year Probability of Fracture; Analysis Restricted to 2^nd^ Generation Parathyroid Hormone Assay Results.

In the multiple variable model predicting vascular events, age, gender, diabetes, obesity, GFR and PTH were significant co-variates (**Table 3**). In this sensitivity analysis, the probability of vascular events was lowest at a baseline PTH of 69 pg/mL (**Figure 3**). In multiple variable models predicting death, age, gender, vascular disease, diabetes, hypertension, hyperlipidemia, obesity, GFR and PTH were significant co-variates (**Table 3**). The probability of death was minimized when baseline PTH was 58 pg/mL (**Figure 3**). The composite outcome of fracture, vascular event or death was lowest at a baseline PTH of 58 pg/mL (**Figure 4**).

**Figure 3:**
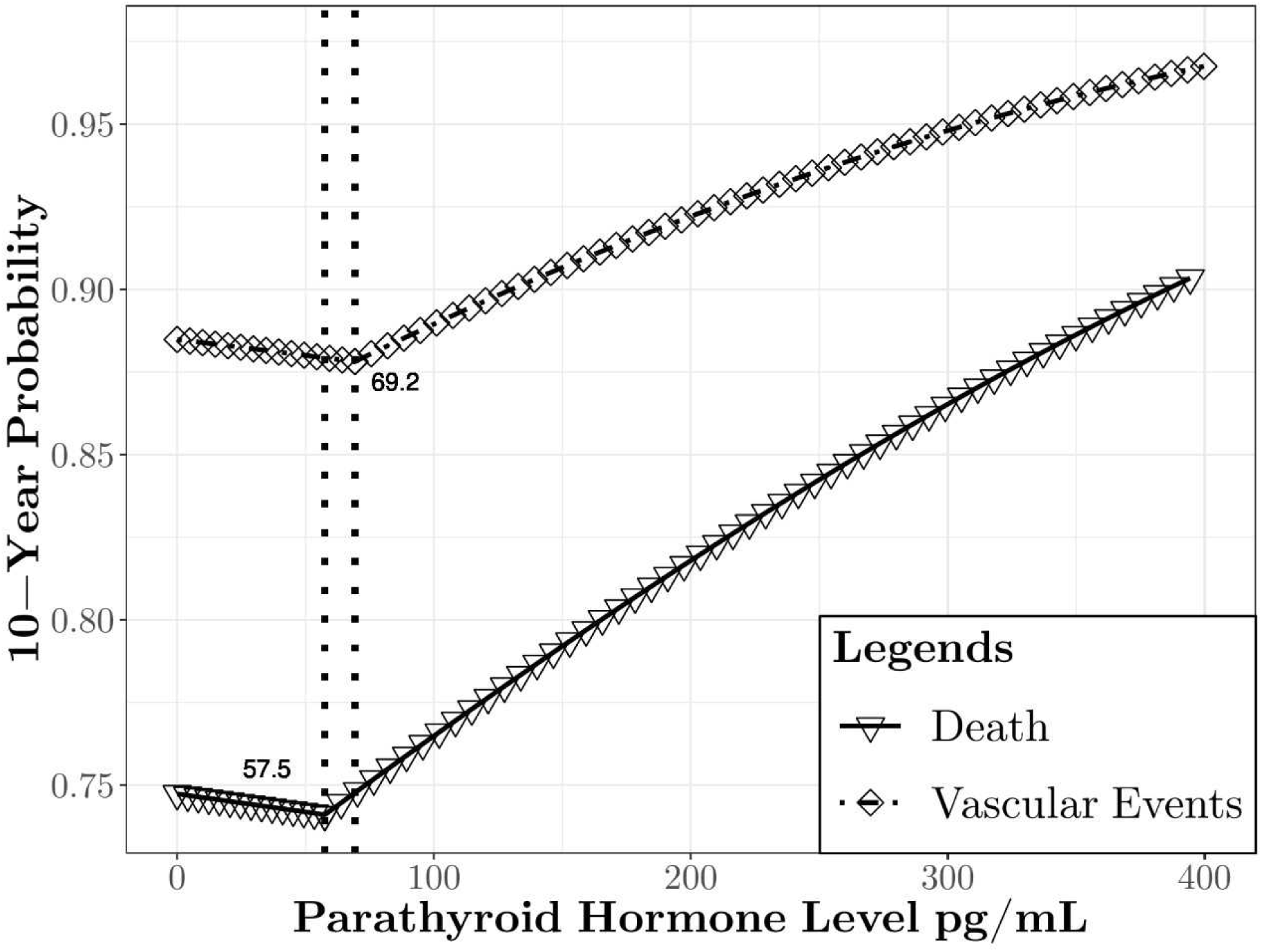
Ten-Year Probability of Vascular Events and Death; Analysis Restricted to 2^nd^ Generation Parathyroid Hormone Assay Results.

**Figure 4:**
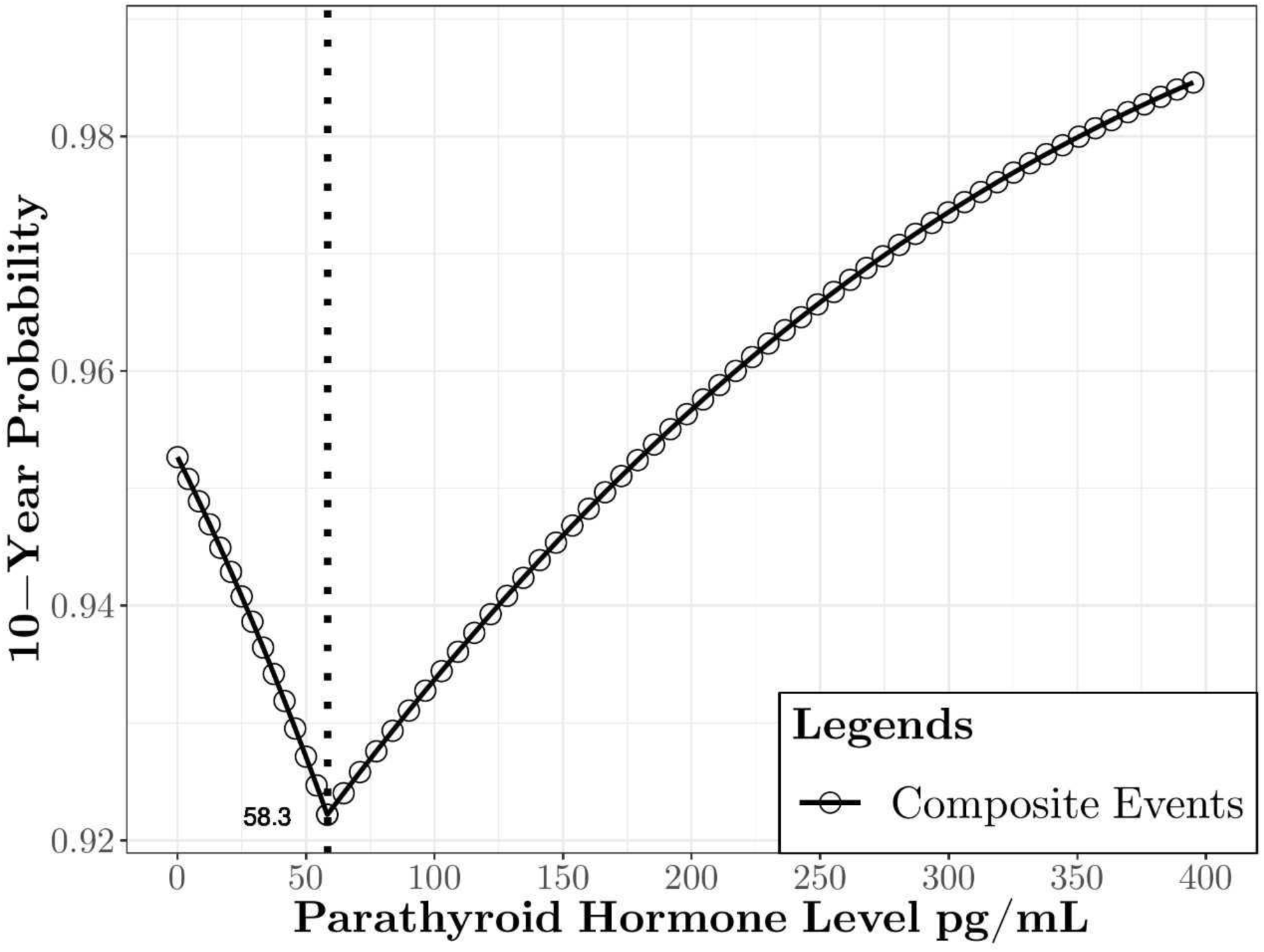
Ten-Year Probability of Fracture; Analysis Restricted to 2^nd^ Generation Parathyroid Hormone Assay Results.

### Probability of Fracture, Vascular Events and Death by stage of CKD

We analyzed the subgroup of 5108 patients to determine whether the optimal PTH cut point differed, based on stage of CKD. In this analysis, we grouped patients into those with an average GFR ≥30 mL/minute (n=3923) and a GFR <30 mL/minute (n=1185) during the baseline year. In these analyses, we found no optimal PTH level that reduced the risk of fracture. In subjects with a GFR ≥30 mL/minute, a PTH of 69 pg/mL was associated with the lowest risk of vascular events. In subjects with a GFR <30 mL/minute, a PTH of 58 pg/mL was associated with lowest risk of death. In the other subgroups, we found no clear cutpoint between PTH and clinical outcomes.

## Discussion

For many years, the National Kidney Foundation recommended treating HPT in CKD patients to achieve specific PTH values (7, 16). The recommendations were based on observational studies linking high PTH to lower BMD and greater risk of vascular events. The Food and Drug Administration approved calcitriol, paricalcitol and doxercalciferol to treat HPT, based on short-term clinical trials demonstrating the ability of these vitamin D compounds to lower PTH values. Thus, clinicians had both guidelines for, and FDA-approved medications to treat, HPT. However in 2017, the National Kidney Foundation recommended against routinely treating HPT (17) due to lack of randomized, placebo-controlled clinical trials proving that such treatment improved patients’ health. We found weak evidence suggested that treating HPT improved bone mineral density (2 trials, 62 subjects) (14, 15) and no clinical trials demonstrating that treatment of HTP reduced vascular events or death. We conducted the current study, in order to evaluate the association between baseline PTH levels and long-term risk of fractures, vascular events and death in a large cohort of patients medically homed at MCHS.

Our main finding was that in patients whose PTH was measured using a 2^nd^ generation assay, fracture risk was lowest at a PTH level of zero, and fracture risk steadily rose in concert with a rising PTH. By contrast, the risk of vascular events and death were lowest when baseline PTH levels were 69 and 58 pg/mL, respectively. The composite endpoint of any adverse event was lowest when PTH equaled 58 pg/mL. In a sensitivity analysis of patients with stage 3 CKD, a PTH of 69 pg/mL was associated with the lowest subsequent risk of vascular events. Likewise in subjects with stage 4 CKD, a PTH of 58 pg/mL was associated with lowest risk of death.

PTH has been criticized as a biomarker in CKD for several reasons. First, biologically inert fragments such as PTH (7-4) accumulate as renal function declines, leading to spurious elevation of the measured PTH. Second, although most laboratories measure PTH using a 2^nd^ generation assays, many such assays are commercially available and each has its “normal” reference range. International standardization is needed, whereby known PTH (1-84) concentrations are measured to confirm consistent results across assays and laboratories. Third, PTH is released in response to ionized serum calcium levels, and thus has diurnal variation influenced by calcium ingestion and intestinal absorption. Reduced PTH variability would be accomplished by measuring PTH in the fasting state. Fourth, magnesium deficiency causes relative hypoparathyroidism and can be challenging to diagnose, as serum magnesium is an insensitive measure of magnesium stores. Finally, biotin supplements can falsely lower PTH values (28) and should be discontinued prior to measuring PTH.

Although PTH may not be an optimal biomarker as discussed above, we found that PTH remained a statistically significant co-variate in the multiple variable models predicting fracture, vascular events and death. The continued use of PTH as a biomarker is supported by our findings. However, several approaches are needed to improve PTH measurement including measurement of PTH in the fasting state, consistent reporting across different commercial assays, and reduced measurement of biologically inert fragments.

We acknowledge that our study has both strengths and limitations. We analyzed a large number of subjects’ electronic health records. We controlled for covariates known to increase the risk of clinical outcomes. However, the nature of this epidemiologic study can only suggest, not prove, a causal relationship between PTH values and clinical outcomes. Additionally, we could not determine whether low PTH values resulted from excess calcium intake, vitamin D therapy or other factors. We had no information on why PTH was measured in subjects, or whether results prompted a specific intervention. Based on census data, our study contained very few non-Caucasian subjects. Finally, we cannot determine whether fractures were due to skeletal fragility, as the circumstances leading fractures were not available via electronic health record search terms.

In conclusion, one in five older adults has CKD and most of these patients develop secondary hyperparathyroidism. While HPT has been linked to lower BMD and greater risk of vascular events, there is no direct evidence from randomized, placebo-controlled trials to confirm that its treatment reduces fractures, vascular events or death. We undertook this study to better understand the relationship between PTH and subsequent risk of fractures, vascular events and death. Our hope was to identify a potential PTH target that would minimize these adverse outcomes, thus guiding the design of randomized controlled clinical trials. Given the limitations of our study, it is important to verify these findings in another database. If we can confirm a proposed target PTH level at which fractures, vascular events and death are reduced, then randomized clinical trials will be needed to verify that treatment of HPT improves the health of CKD patients.

## Author contributions

SG, ZK, DP, DP and PP collected and analyzed the data.

SG created the figures.

KEH designed the study, reviewed analysis of data, drafted and revised the paper.

All authors approved the final version of the manuscript.

## Acknowledgements

Authors received no financial support for the study. We thank Bryan Robeson, Technical Manager at Marshfield Clinic Laboratory for supplying information on the PTH assays used at Marshfield Clinic.

Authors’ disclosures: None

---

**Figure 1 legend**: The figure demonstrates the U-shaped curve of the relationship between baseline parathyroid hormone (PTH) levels and ten-year probability of clinical fractures, vascular events and death. The PTH level at which fractures, vascular events and death was lowest equaled 23 pg/mL, 50 pg/mL and 50 pg/mL, respectively. PTH was measured using a 1^st^, 2^nd^ or 3^rd^ generation assay.

**Figure 2 legend**: The figure demonstrates a linear relationship between baseline parathyroid hormone levels and ten-year probability of clinical fractures. While there was a small inflection point at a PTH value of 101 pg/mL, the hazards of fracture rose steadily with increasing PTH values.

**Figure 3 legend:** The figure demonstrates that hazards of vascular events and death were lowest at PTH values of 69 and 59 ng/mL, respectively.

**Figure 4 legend**: The figure demonstrates that the hazard of any event (fracture, vascular event or death) was lowest at a baseline PTH of 58 pg/mL.

**Appendix Table 1:**
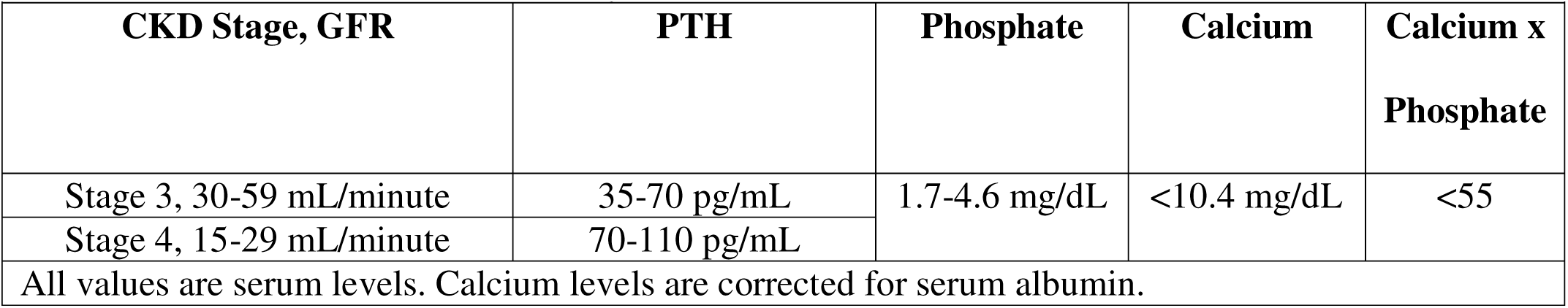
2003 National Kidney Foundation Target Parathyroid Hormone (PTH) Levels

|  |  |  |  |  |
| --- | --- | --- | --- | --- |
| CKD Stage, GFR | PTH | Phosphate | Calcium | Calcium x<br>Phosphate |
| Stage 3, 30-59 mL/minute | 35-70 pg/mL | 1.7-4.6 mg/dL | <10.4 mg/dL | <55 |
| Stage 4, 15-29 mL/minute | 70-110 pg/mL |  |  |  |
| All values are serum levels. Calcium levels are corrected for serum albumin. |  |  |  |  |

**Appendix Table 2:**
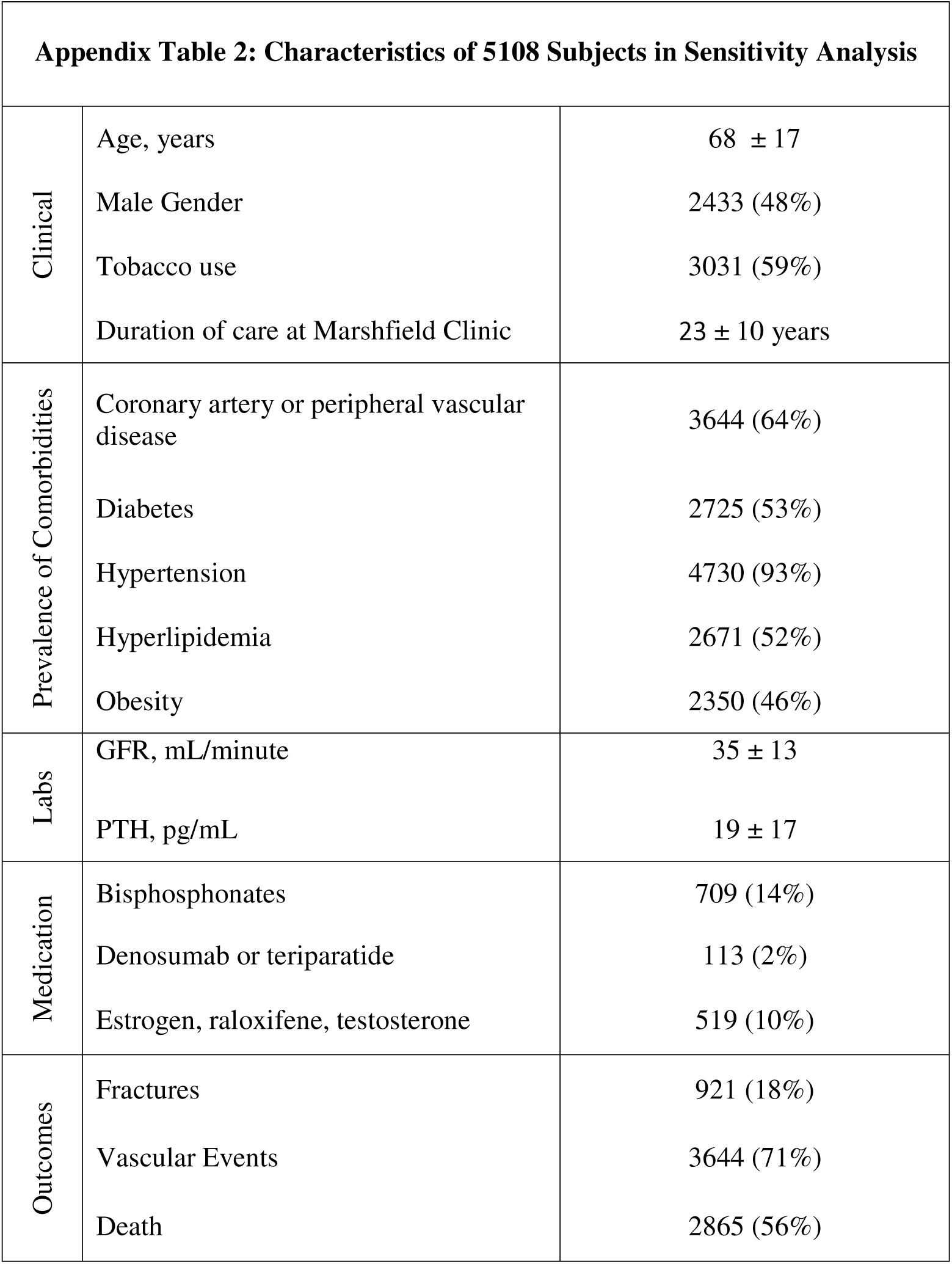
Characteristics of 5108 Subjects in Sensitivity Analysis

|  |  |  |
| --- | --- | --- |
| Clinical | Age, years | 68 ± 17 |
|  | Male Gender | 2433 (48%) |
|  | Tobacco use | 3031 (59%) |
|  | Duration of care at Marshfield Clinic | 23 ± 10 years |
| Prevalence of Comorbidities | Coronary artery or peripheral vascular disease | 3644 (64%) |
|  | Diabetes | 2725 (53%) |
|  | Hypertension | 4730 (93%) |
|  | Hyperlipidemia | 2671 (52%) |
|  | Obesity | 2350 (46%) |
| Labs | GFR, mL/minute | 35 ± 13 |
|  | PTH, pg/mL | 19 ± 17 |
| Medication | Bisphosphonates | 709 (14%) |
|  | Denosumab or teriparatide | 113 (2%) |
|  | Estrogen, raloxifene, testosterone | 519 (10%) |
| Outcomes | Fractures | 921 (18%) |
|  | Vascular Events | 3644 (71%) |
|  | Death | 2865 (56%) |

